# Progress in elementary school reading linked to growth of cortical responses to familiar letter combinations within visual word forms

**DOI:** 10.1101/2022.11.08.515712

**Authors:** Fang Wang, Blair Kaneshiro, Elizabeth Y. Toomarian, Radhika S. Gosavi, Lindsey R. Hasak, Suanna Moron, Quynh Trang H. Nguyen, Anthony M. Norcia, Bruce D. McCandliss

## Abstract

Learning to read depends on the ability to extract precise details of letter combinations, which convey critical information that distinguishes tens of thousands of visual word forms. To support fluent reading skill, one crucial neural developmental process is one’s brain sensitivity to statistical constraints inherent in combining letters into visual word forms. To test this idea in early readers, we tracked the impact of two years of schooling on within-subject longitudinal changes in cortical responses to different properties of words (coarse tuning for print, and fine tuning to visual word forms and whole word representations) and their growth in reading skill. Three stimulus contrasts—words versus pseudofonts, words versus pseudowords, pseudowords versus nonwords—were presented while high-density EEG Steady-State Visual Evoked Potentials (SSVEPs, n=31) were recorded. Internalization of abstract visual word form structures over two years of reading experience resulted in a near doubling of SSVEP amplitude, with increasing left lateralization. Longitudinal changes in brain responses to such word form structural information were linked to the growth in reading, especially in rapid automatic naming for letters. No such changes were observed for whole word representation processing and coarse tuning for print. Collectively, these findings indicate that sensitivity to visual word form structure develops rapidly through exposure to print and is linked to growth in reading skill.

**Research Highlights:**

- Longitudinal changes in cognitive responses to coarse print tuning, visual word from structure, and whole word representation were examined in early readers.
- Visual word form structure processing demonstrated striking patterns of growth with nearly doubled in EEG amplitude and increased left lateralization.
- Longitudinal changes in brain responses to visual word form structural information were linked to the growth in rapid automatic naming for letters.
- No longitudinal changes were observed for whole word representation processing and coarse tuning for print.

## 1 Introduction

Visual word recognition is a rapid yet complex process involving multiple dimensions of analysis, including the specific shape of each letter form, the rules of letter combination within a word form, and whole word representation. Given this complexity, it has long been an ambition of developmental and educational cognitive neuroscience research to clarify how emerging expertise at different levels of word processing is connected with changes in visual reading circuits and how these neural changes influence reading skills. Such work can help elucidate the neural basis of individual differences in reading acquisition and reading skills, and also help demonstrate within-individual changes due to specific educational activities.

A large body of literature (Bentin et al., 1999; Brem et al., 2005; Eberhard-Moscicka et al., 2015; Maurer et al., 2006; Wang & Maurer, 2017) has investigated visual word recognition by contrasting responses to words versus visual controls (e.g., pseudofonts, strings combined with artificial character set font). Taken together, studies have consistently reported brain sensitivity to words compared to pseudofonts, a finding that is referred to as “coarse neural tuning” for print (Maurer et al., 2006). During development, coarse neural tuning starts to emerge when children begin to read, following an inverted U-curve with an initial increase (van de Walle de Ghelcke et al., 2021) and then a later decrease starting in second grade (Maurer et al., 2006, 2010).

The often-used coarse neural tuning contrast (words versus pseudofonts) is limited, however, in its connection to theory-driven insights into word recognition development, which stress the importance of multiple levels of word reading expertise (Carreiras et al., 2014). This limitation leads to ambiguity in recovering the unique contributions of specific properties of words to developmental changes in reading and brain circuitry. Thus it is an open question whether coarse neural tuning reflects merely the aggregation of increasing word specific knowledge, or instead reflects insights into word form structures, perhaps as they map to phonological patterns (Maurer & McCandliss, 2007).

Event-related potential (ERP) studies have attempted yet failed to isolate different levels of visual word processing in early readers (Eberhard-Moscicka et al., 2015; Zhao et al., 2014). For instance, Zhao and colleagues found brain sensitivity to visual word form structure only in 7-year-old children with high reading fluency, but failed to reveal sensitivity to whole visual word knowledge (Zhao et al., 2014). At the same time, steady-state visual evoked potential (SSVEP) paradigms have been increasingly used, as this approach rapidly measures discrimination responses with high signal-to-noise ratio (SNR) in only a few minutes of stimulation (Norcia et al., 2015) and has high test-retest reliability (Dzhelyova et al., 2019). Despite these advantages, SSVEP studies on word reading have not yet successfully differentiated effects of learning at the whole word level from effects of word form structure learning in early readers (i.e., kindergartners in Lochy et al. (2016) or first and second graders in van de Walle de Ghelcke et al. (2021)). More importantly, no study so far has systematically tracked the developmental trajectories and neural dynamics of multiple levels of word reading expertise in early reading acquisition.

The current study aims to address this gap by isolating different levels of word reading expertise that emerge over the course of reading development, and by better connecting brain signals related to these different levels of word recognition with early reading fluency and reading growth.

In our recent SSVEP study (Wang et al., 2022), we isolated different levels of word-related information using multiple well-controlled contrasts and an adjusted SSVEP paradigm. Typically, SSVEP paradigms involve presentation of two categories of stimuli that differ in a particular aspect (e.g., words versus pseudofonts) at two distinct, experimentally defined periodic rates. Previous SSVEP studies have used this “base/deviant” approach at faster stimulation rates, wherein deviant stimuli were embedded within a sub-multiple of the base rate that is greater than two, for example, 1.2 Hz deviant and 6 Hz base rates. We used an adjusted paradigm—which we refer to as “image alternation” mode—involving slower stimulation rates, where image exemplars from two categories of stimuli alternate at slower 1 Hz alternation and 2 Hz base rates. This slower image alternation mode has been shown to elicit responses with a higher SNR (Yeatman & Norcia, 2016; Barzegaran & Norcia, 2020; Wang et al., 2021) than base/deviant approaches and may thus be more suitable for capturing reading-related brain signals in children (Wang et al., 2022).

We used three stimulus contrasts. First, the commonly used words–pseudofonts contrast was included to probe coarse neural tuning. Second, we used a pseudowords–nonwords contrast to investigate responses specific to orthographic structures within visual word forms while controlling for visual familiarity and whole word representation. In this contrast, well-structured orthographically reasonable letter combinations (pseudowords) were alternated with letter strings that violate orthographic constraints (nonwords). Notably, to avoid potential confounding of consonant-vowel distributions, orthographically illegal nonwords were created by reordering letters from consonant-vowel-consonant (CVC) pseudowords instead of pure consonant strings used in previous contrasts (Zhao et al., 2014). Finally, responses to whole visual words was examined through a words–pseudowords contrast, which serves to contrast information on the level of a word unit while controlling for letter and letter combination structure experience.

In all, the stimulus contrasts and stimulation rates in the present study are those used in our recent non-longitudinal SSVEP study (Wang et al., 2022). Additionally and most importantly, longitudinal EEG data were recorded over a two-year interval in order to (1) track the developmental profile of different levels of word information, and (2) clarify to what extent sensitivity to visual word form structure and/or sensitivity to whole word knowledge relates to reading fluency and reading growth.

## 2 Methods

### 2.1 Participant Sample

Thirty-three English-speaking children, with normal or corrected-to-normal vision and no reading disabilities, completed behavioral and EEG sessions at two time points. Samples were collected during October through November, 2019 (T1^1^), and again two years later (T2). Two participants were excluded due to data quality issues, resulting in *N* = 31 participants (16 males, 15 females), whose EEG data were analyzed. The T1 sample included 10 kindergarteners, 12 first graders, and 9 second graders (*m* = 6.78 years, *s* = 0.81 years), and correspondingly 10 second graders, 12 third graders, and 9 fourth graders at T2 (mean = 8.78 years, *s* = 0.79 years).

### 2.2 Behavioral Assessment

Behavioral assessments were administered in separate sessions on average 7.97 (*s* = 4.97) days and 4.58 (*s* = 7.21) days either before or after the EEG session for T1 and T2, respectively. All children were tested on two sub-tests of the Comprehensive Test of Phonological Processing, Second Edition (CTOPP-II, Wagner et al. (2013))—Rapid Automatized Naming (RAN) of colors and letters—for their phonological awareness and rapid naming abilities. In addition, the Sight Word Efficiency (SWE) and the Phonemic Decoding Efficiency (PDE) subtests of the Test of Word Reading Efficiency, Second Edition, (TOWRE-II, Torgesen et al. (2012)) were used to measure efficiency of sight word recognition and phonemic decoding ability. We used the letter-word identification subset of Woodcock-Johnson Tests of Achievement, Fourth Edition (WJ-IV; Schrank et al. (2014)) to assess word decoding ability. Finally, the handedness of each participant was defined by the Edinburgh Handedness Inventory (Oldfield, 1971). Participant demographics and results of behavioral assessments are summarized in Table 1.

**Table 1:** Subject characteristics and behavioral assessments at T1 and T2. Values are mean(SD). TOWRE: Number of real words and pronounceable nonwords read in 45 seconds. RAN: Time (ms) used to quickly and accurately name all stimuli (e.g., letters or colors) on a test form. WJ: Number of correctly named letters and words until getting 6 in a row wrong. T1: First testing time; T2: Second testing time (two years later).

|  | Group 1 | Group 2 | Group 3 |
| --- | --- | --- | --- |
| Sex (female/male) | 3/7 | 6/6 | 6/3 |
| Handedness (right/left) | 9/1 | 10/2 | 8/1 |
| **T1** | Kindergarten | Grade 1 | Grade 2 |
| Age in years | 5.88($\pm 0.42$) | 6.81($\pm 0.31$) | 7.74($\pm 0.24$) |
| Test of word reading efficiency (TOWRE) | 10.55($\pm 9.32$) | 63.33($\pm 28.07$) | 79.22($\pm 22.73$) |
| Rapid automatic naming of colors (RANcolor) | 42.80($\pm 9.02$) | 37.50($\pm 7.66$) | 36.89($\pm 9.23$) |
| Rapid automatic naming of letters (RANletter) | 45.11($\pm 15.00$) | 25.33($\pm 6.34$) | 22.33($\pm 2.78$) |
| Word decoding ability (WJ) | 15.50($\pm 5.38$) | 43.83($\pm 14.02$) | 51.44($\pm 8.56$) |
| **T2** | Grade 2 | Grade 3 | Grade 4 |
| Age in years | 7.90($\pm 0.38$) | 8.80($\pm 0.34$) | 9.73($\pm 0.23$) |
| Test of word reading efficiency (TOWRE) | 79.30($\pm 21.11$) | 100.42($\pm 13.46$) | 104.11($\pm 19.41$) |
| Rapid automatic naming of colors (RANcolor) | 30.20($\pm 4.96$) | 31.42($\pm 5.99$) | 27.63($\pm 7.40$) |
| Rapid automatic naming of letters (RANletter) | 21.09($\pm 3.20$) | 19.03($\pm 3.89$) | 18.49($\pm 4.27$) |
| Word decoding ability (WJ) | 54.11($\pm 7.77$) | 62.33($\pm 5.10$) | 62.22($\pm 3.80$) |

### 2.3 Stimuli and Experimental Paradigm

Four types of stimuli were used: Words, pseudofonts, pseudowords, and nonwords. Words were high frequency (average: 2110 per million, range: 629–5851 per million) three-letter CVC English words rendered in Courier New font. Pseudofonts were word stimuli presented in the Brussels Artificial Character Set font (BACS-2, Vidal & Chetail (2017)), which provide well-matched visual controls of Courier New letters. Pseudowords were generated on an item-by-item basis by semi-randomly rearranging letters of words stimuli while retaining a CVC structure, rendering them still pronounceable with well-matched bigram frequencies (*t*(30) = 0.26*, p* = 0.79) and orthographic neighborhood sizes (*t*(30) = 0.79*, p* = 0.43) with words. Finally, nonwords stimuli were also built on an item-by-item basis by semirandomly shuffling letters across pseudowords to produce unpronounceable exemplars not following English orthographic and phonological rules. Bigram frequencies and orthographic neighborhood sizes of nonwords and pseudowords differed significantly (both *t*(30) *>* 6.26, both *p <* 0.001). In all, we prepared 32 high-frequency W, 16 PF, 32 PW and 16 NW, for a total of 96 stimulus exemplars. All stimuli images, spanning 9.4 (horizontal) by 2.5 (vertical) degrees of visual angle, were presented in black font on a gray background at the center of the screen. Stimuli images and background contrast was set at 95%.

Using these four categories of stimuli, three different contrast conditions were investigated: Coarse print tuning (words–pseudofonts alternation), whole word representation processing (words–pseudowords alternation), and sensitivity to visual word form structure (pseudowords–nonwords alternation). The three conditions were presented in the aforementioned fixed order (see Limitations). To avoid potential learning effects from stimulus repetition in SSVEP studies (De Rosa et al., 2022), the 16 words used in the words–pseudofonts condition differ from those words used in words–pseudowords; the 16 pseudowords in words–pseudowords differ from those in pseudowords–nonwords. Each condition started with a blank screen shown for a random interval of 1500–2500 ms. Then, a given image was presented for 500 ms, with a complete cycle of stimulus alternation lasting 1000 ms; these periods denote the 2-Hz *base frequency* (i.e., 2 items per second) and 1-Hz *alternate frequency* (i.e., one item/alternate per second), respectively. Stimulus conditions and examples are shown in Figure 1. Participants performed a repetition detection task, pressing a button on a response pad with their preferred hand after detection of a stimulus presented three times consecutively (Figure 2).

**Figure 1:**
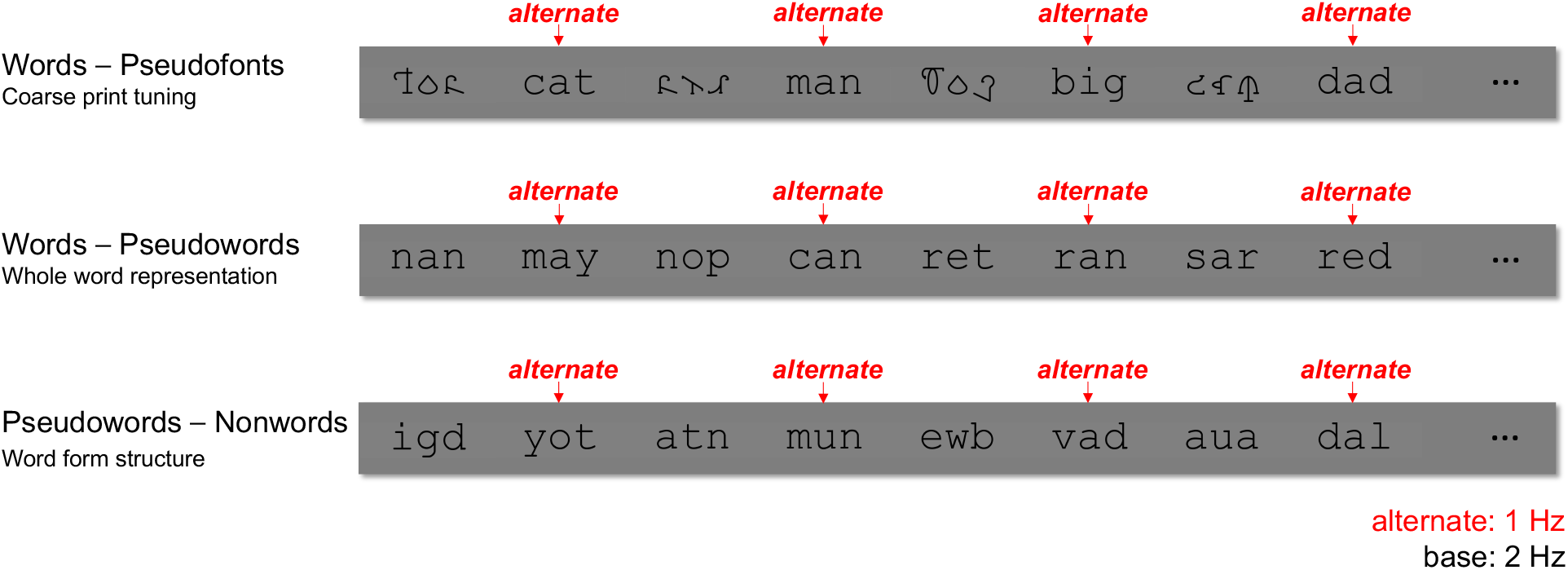
Experimental Design. Examples of stimuli presented in the experiment. 1 Hz alternates were embedded within a 2 Hz base stream in all three conditions. The first condition assessed coarse print tuning with words alternating with pseudofonts (W–PF). The second condition assessed lexical fine tuning with words alternating with orthographically legal pseudowords (W–PW). The third condition assessed sublexical fine tuning with orthographically legal pseudowords alternating with orthographically illegal nonwords (PW–NW). All contrasts were presented centered on the screen.

**Figure 2:**
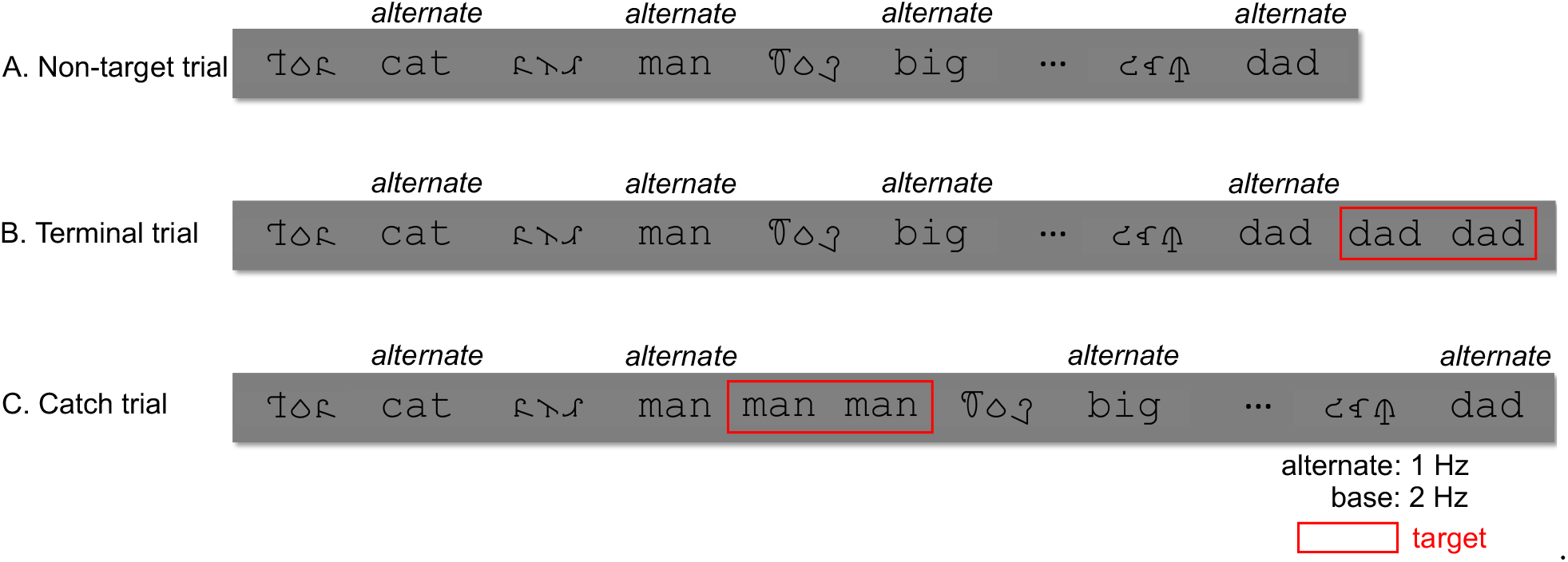
Different types of trials in words–pseudofonts contrast. Twelve trials of prerandomized sequences were presented for each condition (words–pseudofonts contrast shown here as an example): Four non-target trials (A), four terminal trials (B), and four catch trials (C). EEG data corresponding to the four catch trials were excluded from analysis due to excessive responserelated movements during recording. For the eight trials included per participant, the first and last 1-second epochs of each 12-second trial were excluded to avoid transient responses associated with ocular artifacts which occurred more at beginnings and endings of trials

Twelve successive 1-second epochs (each containing 2 stimuli: 1 *alternative* stimulus, 1 control stimulus) comprised one trial. For each condition, 12 trials were presented in pre-randomized sequences. Four “non-target” trials, four “terminal” trials, and four “catch” trials depending on whether and when repetition targets appeared. Specifically, non-target trials contained no repeated stimuli (Figure 2A); in terminal trials, repeated stimuli appeared at the end (Figure 2B); while repeated stimuli randomly appeared elsewhere during catch trials (Figure 2C). Only one target appeared in each terminal and catch trial. Feedback on task performance was provided after the end of each trial. Participants were given breaks as needed between trials.

Because EEG data from catch trials contained movement artifacts related to the key press, EEG data corresponding only to the four non-target trials and four terminal trials were analyzed. For these 8 trials included per participant, the first and last 1-second epochs of each 12-second trial were excluded to avoid transient responses associated with ocular artifacts which occurred more at beginnings and endings of trials. In all, 10 epochs (i.e., seconds) of data from each of 8 trials per participant were analyzed.

### 2.4 Experimental Procedure and Data Acquisition

Participants sat in a dimly lit room 1 m away from a computer screen. Prior to EEG recording, participants were instructed on the repetition detection task and performed a brief practice session with corrective feedback. EEG sessions took approximately 45–50 minutes, including setup and between-trial breaks. As compensation, each child received a small token gift after the study.

128-sensor EEG (Electrical Geodesics, EGI NA 400 amplifier) were recorded against Cz reference, sampled at 500 Hz. Impedances were kept below 50 kΩ. Recorded data were digitally low-pass (50 Hz) and high-pass (0.3 Hz) filtered offline using Net Station Waveform Tools. Filtered data were then imported into in-house signal processing software for preprocessing.

### 2.5 EEG Preprocessing and Analysis

During preprocessing, EEG data were first re-sampled to 420 Hz. This was done to ensure an integer number (herein 7) of time samples per video frame given the frame rate of 60 Hz, as well as integer numbers of frames (i.e., 60 frames per 1 Hz cycle and 30 frames per 2 Hz cycle) per stimulus cycle given the current stimulation frequencies of 1 Hz and 2 Hz. Then, sensors were interpolated with the average of data from six neighboring sensors if more than 15 % of samples exceeded a ± 60 *µV* amplitude threshold. The continuous EEG data were then re-referenced to average reference (Lehmann & Skrandies, 1980) and segmented into 1-second epochs. Epochs with more than 10 % of time samples exceeding a ± 60 *µV* noise threshold, or with any time sample exceeding an artifact threshold of (± 70 *µV*) (reflecting e.g., eye blinks, eye movements or body movements), were rejected from further analyses.

Recursive Least Squares (RLS) filters were then used to filter the epoched EEG signal in the time domain (Tang & Norcia, 1995). The filters were tuned to each of the analysis frequencies (i.e., alternate frequency 1 Hz, base frequency 2 Hz, and their harmonics) and converted to complex amplitude values by means of the Fourier transform. Complex-valued RLS outputs were decomposed into real and imaginary Fourier coefficients as input for subsequent analyses.

#### 2.5.1 Reliable Components Analysis (RCA)

Instead of analyzing periodic responses from several pre-selected (literature- or SNR-based) sensors that may lead to a bias in reporting false positives (Kilner, 2013) and fail to separately describe activities underlying multiple cortical sources (Wang et al., 2021), we spatially filtered the sensor-space data using RCA. RCA has been shown to capture and isolate different neural processes arising from different underlying sources (Dmochowski et al., 2012), and topographies of corresponding spatial filters can strongly resemble lead fields generating observed SSVEPs (Dmochowski et al., 2015).

RCA is a matrix decomposition technique, which decomposes the entire 128-sensor array into a set of reliable components that maximizes between-trial covariance relative to within-trial covariance (Dmochowski et al., 2012, 2015). Given a sensor-by-feature (i.e., real and imaginary Fourier coefficients at selected frequencies of interest) EEG data matrix, RCA computes optimal weightings of sensors (i.e., linear spatial filters), through which the sensor-by-feature matrices transformed to component-by-feature matrices, with each component representing a linear combination of sensor (Dmochowski et al., 2015).

The resulting weight vectors (spatial filters) are vectors of length *N_sensor_* which represent linear weightings of sensors (i.e., linear spatial filters). Sensor-space EEG data can then be projected through (i.e., multiplied by) these weight vectors to transform the spatial dimension of the data from sensors to spatial components, where the number of components is typically much smaller than the number of sensors. The spatial filtering process of RCA is conceptually similar to process of deriving linear spatial components using Principal Components Analysis (PCA) in that both involve eigenvalue decompositions returning multiple components sorted in descending order of criteria explained; however, where PCA optimizes sensor weightings to maximize variance explained within one data matrix, RCA computes sets of weights to maximize covariance across pairs of data matrices (trials). Moreover, in contrast to PCA, the spatial filters computed by RCA are not constrained to be orthogonal (Dmochowski et al., 2012). Technical details on the RCA technique are provided by Dmochowski et al. (2012, 2015).

RCA produces three outputs that are of interest in our present analyses. First, the weight vectors themselves, which are the eigenvectors of the RCA calculation, can be visualized as scalp topographies using a forward-model projection (Parra et al., 2005). These visualizations are thought to reflect the propagation of synchronized activity captured by the weight vectors onto the scalp (Parra et al., 2005). Next, each weight vector (eigenvector) is accompanied by an eigenvalue, which we refer to as the coefficient of a given component. These coefficients give a measure of optimized across-trials correlation for each component. Finally, the component-space data (i.e., the spatially filtered sensor-space data) undergo subsequent analysis and visualization.

#### 2.5.2 RCA Calculations

In order to test whether low-level visual features were well matched across conditions, we computed RCA on base frequency and its harmonics. Specifically, we computed RCA over the first five harmonics of the base (2 Hz, 4 Hz, 6 Hz, 8 Hz, and 10 Hz). As done in a previous analysis of a superset of T1 data (Wang et al., 2022), these calculations were performed on all conditions together to enable direct quantitative comparisons of responses to low-level visual stimulus features in a shared component space. However, we computed RCA for T1 and T2 separately in order to investigate potential developmental changes of such responses. To assess the processing difference between alternate and control stimuli, we computed RCA over the first five odd harmonics of alternate (1 Hz, 3 Hz, 5 Hz, 7 Hz, and 9 Hz), excluding even/base harmonics (2 Hz, 4 Hz, 6 Hz, 8 Hz, and 10 Hz). We computed RCA separately for each condition and testing time point to investigate different neural mechanisms underpinning multiple levels of information processing that were evoked by different stimulus contrast conditions over developmental stages.

#### 2.5.3 Statistical Analyses of RC Data

Statistical significance of the coefficients (eigenvalues) of RCA components was assessed using permutation test. As in Wang et al. (2022), we generated surrogate versions of the sensor-space data: For every 1-second epoch of a 10-second trial (accounting for within-trial, across-epoch correlations introduced by RLS filtering) the phases of the data were rotated by a random angle, independently for each harmonic. We computed RCA over 500 such surrogate versions of the data and treated the resulting distributions of RCA coefficients as null distributions for computing p-values of the observed (intact) coefficients. These p-values were then corrected for multiple comparisons using FDR (Benjamini & Yekutieli, 2001). The significance of coefficients for *alternate* RCA calculations were tested separately for each condition, given RCA was computed separately for each condition in order to investigate different neural mechanisms underpinning multiple levels of word information processing. In contrast, the significance of coefficients for base RCA calculations was tested on three conditions together, as the calculation of spatial filters on three conditions together enables direct quantitative comparisons of responses to low-level visual stimulus features in a shared component space (Wang et al., 2022).

For each RC, a 1-second epoch of frequency-domain component-space data contained 10 data points (5 harmonics times 2 real and imaginary coefficients). Component-space data were first averaged across 1-second epochs on a per-participant basis. Following this, statistical analyses were performed across distributions of participants. We performed Hotelling’s two-sample t^2^ tests (Victor & Mast, 1991) on distributions of real and imaginary Fourier coefficients on a per-harmonic, per-component basis to identify responses that differed significantly from zero in the complex plane, correcting for multiple comparisons using False Discovery Rate (FDR, Benjamini & Yekutieli (2001)).

To identify and compare overall response amplitude *across* multiple significant harmonics, we combined harmonic response amplitude using the root sum of squares (henceforth referred to as *RSS* projected amplitude), that is, the square root of the summed squared amplitudes at multiple harmonics (Tlumak et al., 2011; Appelbaum et al., 2010).

For alternate harmonics results (RCA trained on each condition separately), we first computed a two-way repeated-measures ANOVA with within-subjects factors of condition and testing time point on response amplitude (RSS amplitude) across three stimulus contrasts. Then, we performed paired-sample, two-tailed t tests on *RSS* projected amplitudes across testing time points for RC separately in each condition; for words–pseudowords and pseudowords–nonwords, where only the first harmonic (1 Hz) was significant, we further performed paired-sample t tests (two tailed) on projected amplitude at 1 Hz. For base harmonics results (RCA trained on three conditions together), we performed a two-way repeated-measures ANOVA (within-subjects factors of condition and testing time point) on *RSS* projected amplitudes for each RC separately.

We used the Circular Statistics toolbox (Berens et al., 2009) to compare distributions of RC phases for harmonics with significant responses at both testing time points. For alternate harmonics, we computed circular t tests at a given harmonic between two testing points; here, words–pseudofonts results were corrected for 3 comparisons, while words–pseudowords and pseudowords-nonwords involved no multiple comparisons as only the first harmonic was significant. For base harmonics, using the Circular Statistics Toolbox (Berens et al., 2009), we computed circular ANOVA across three conditions at each harmonic within time point, followed by pairwise circular t tests; results were FDR-corrected across 5 comparisons (5 significant harmonics).

#### 2.5.4 Visualization of RCA Data

In visualizing the EEG results, we first present scalp topographies (i.e., forward-model projection of the spatial filters) of reliable components. Second, we present mean responses of projected data as vectors in the 2D complex plane: The vector length represents the response amplitude and the angle of the vector relative to 0 degrees (counterclockwise from the 3 o’clock direction) represents the phase. Error ellipses around the vectors are the standard errors of the mean (*SEM*). Third, we present amplitudes (*µV*) at each harmonic for each component in bar plots, with statistically significant responses indicated with asterisks, as determined by adjusted *p_FDR_* values of Hotelling’s two-sample t^2^ tests of combined real and imaginary coefficients.

### 2.6 Analysis of brain-behavior relationships

We explored possible brain-behavior relationships using *RSS* projected amplitudes from alternate RCA output for words–pseudofonts and 1 Hz amplitude for words–pseudowords and pseudowords–nonwords in conjunction with reading scores. To test the association between brain signals and reading performance at different developmental stages, we performed linear regressions of response amplitude with each of the reading scores (i.e., RANcolor, RANletter, TOWRE, and WJ), separately for T1 and T2. Next, to relate the development of brain responses to changes in reading performance, we computed linear regression of the change in response amplitude from T1 to T2 (*RSS_T_*_2_ *−RSS_T_*_1_ for words–pseudofonts, amplitude at 1 Hz*_T_*_2_ *−* amplitude at 1 Hz*_T_*_1_ for words–pseudowords and pseudowords–nonwords), with corresponding reading performance improvements (i.e., RANcolor*_T_*_2_ *−* RANcolor*_T_*_1_, RANletter*_T_*_2_ *−* RANletter*_T_*_1_, TOWRE*_T_*_2_ *−* TOWRE*_T_*_1_, and WJ*_T_*_2_ *−* WJ*_T_*_1_). Outliers determined by Cook’s Distance (Cook, 1977) based on the regression model were removed if they exceeded the 4*/n* threshold (*n* total data points).

### 2.7 Behavioral Analysis

For the repetition detection task performed during EEG sessions, we computed *d’* based on the z-transformed probabilities of hits and false alarms (Macmillan & Creelman, 2004). A two-way repeated-measures ANOVA with within-subjects factors of condition and testing time point was computed on *d’* across conditions and time points.

## 3 Results

### 3.1 Alternate RCA Results

Reliable component (RC) coefficients, which indicate the extent of across-trials covariance explained by a given component, dropped steeply after the first RC (see supplement, Figure S2). We therefore focused our alternate RCA results solely on the maximally correlated component, RC1. A two-way repeated-measures ANOVA showed a significant main effect of condition (F(2,60) = 56.80, p<0.001) and a significant interaction effect between condition and testing time point (F(2,60) = 6.77, p<0.01). No significant main effect of testing time point was found (F(1,30) = 0.08, p=0.78).

#### 3.1.1 Development of sensitivity to word form structure (pseudowords–nonwords)

For responses to pseudowords–nonwords, the contrast designed to probe visual word form structure processing, RC1 is maximal over occipital electrodes on both hemispheres at the first testing time point, T1 (Figure 3A). Responses are statistically significant at the first two harmonics 1 Hz and 3 Hz (Hotelling’s two-sample t^2^ test, *p_FDR_ <* 0.01, corrected for 10 comparisons, Figure 3C). At the second testing time point, T2, the activation is more left lateralized (Figure 3B) with statistically significant responses at only the first harmonic, 1 Hz (*p_FDR_ <* 0.001, corrected for 10 comparisons, Figure 3C). Developments of lateralizaion of brain responses at T1 and T2 are shown in Figure S5B.

**Figure 3:**
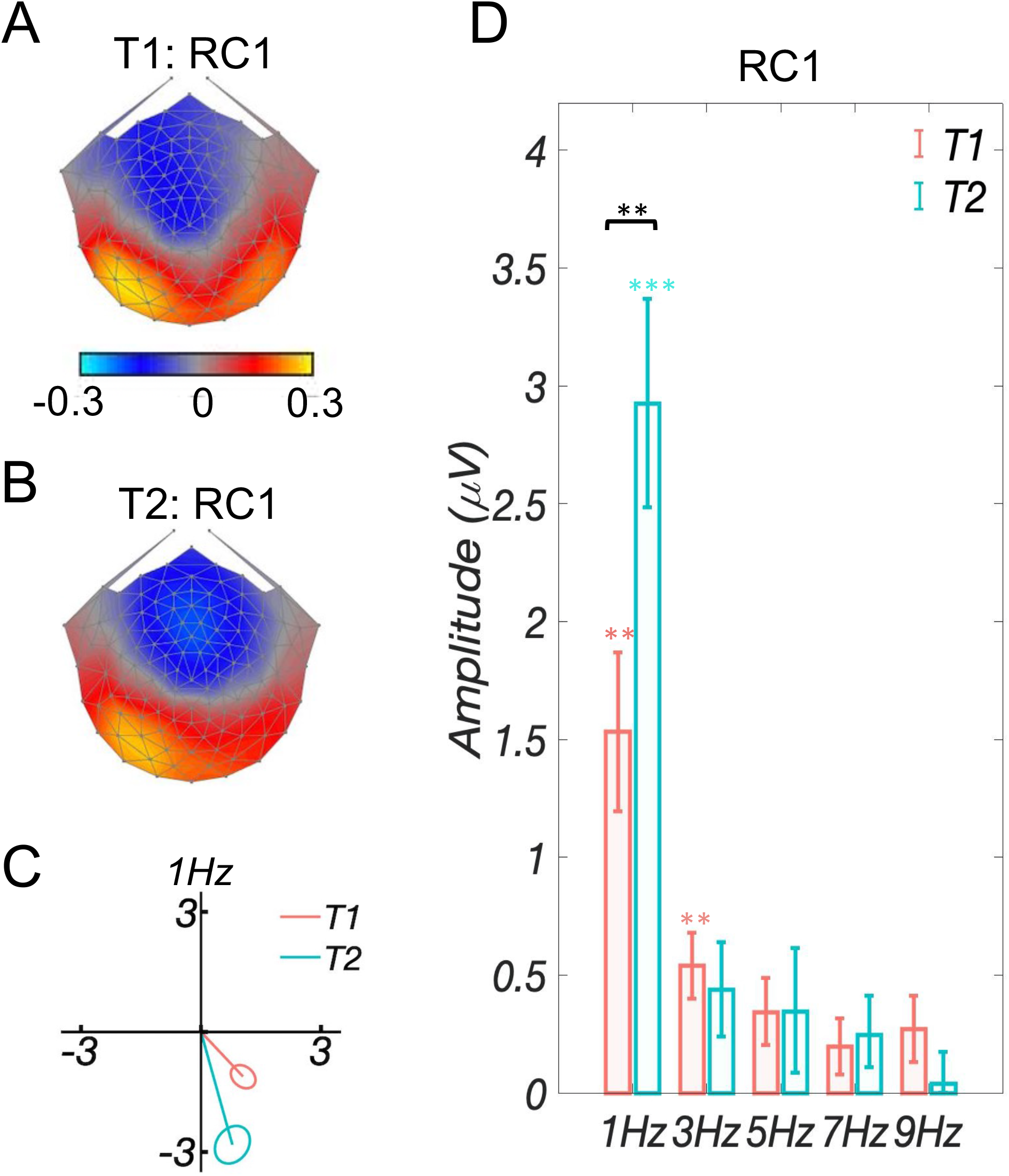
Alternate Analysis: Developmental changes of visual word form structure tuning (pseudowords–nonwords) in early readers. A: Topographic visualization of the spatial filter (bilateral OT) for the maximally reliable component (RC1) at T1; B: Topographic visualization of the spatial filter (left OT) for RC1 at T2; C: Response data at the first harmonic (1 Hz, the only significant harmonic at both testing time points) presented in the complex plane: Amplitude information is represented as the length of the vector, and phase information in the angle of the vector relative to 0 degree (counterclockwise from 3 o’clock direction), ellipses indicate standard error of the mean (*SEM*). No significant phase difference (*p* = 0.55) was found between T1 and T2; D: Projected amplitude for each harmonic at T1 (orange) and T2 (blue), respectively. The first two harmonics were significant at T1, while only the first harmonic was significant at T2. A significant difference (*p <* 0.01) was found at 1 Hz between T1 and T2. No significant differences were revealed at other harmonics. **: *p_F DR_ <* 0.01, ***: *p_F DR_ <* 0.001.

In order to investigate changes in response amplitude across testing time points, we computed paired sample t-tests (two tailed) of amplitude. We considered only the first two harmonics (1 Hz and 3 Hz), for which responses were significant at at least one testing time point. Paired t-test (two tailed) of RC1 amplitudes at 1 Hz showed that amplitudes at T2 are significantly higher than (nearly double) those at T1 (*t*(1, 30) = 2.74*, p_FDR_ <* 0.01, corrected for two comparisons), while no significant difference between testing time points was observed at 3 Hz (*t*(1, 30) = 0.71*, p_FDR_* = 0.48, corrected for two comparisons). There was no significant difference in phase at 1 Hz (the only significant harmonic at both testing time points) between the testing time points (circular t-test; *p* = 0.55).

#### 3.1.2 Development of whole word representation processing (words–pseudowords)

The words–pseudowords contrast was designed to detect whole word representation processing. We first calculated RCA on each testing time point separately, as was done for the other two contrasts. Since EEG responses to this contrast were relatively weak, we further pooled the sensor-space data across testing time points to calculate the RCA filters.

As shown in Figure 4A&B, the topographies of RC1 trained on T1 and T2 separately include activation over occipital and temporal electrodes, with relatively weak and noisy responses. The topography of RC1 trained on the two time points together includes activation at more anterior left vOT electrodes (Figure 4C). Projected data were statistically significant at 1 Hz (*p_FDR_ <* 0.01) and 5 Hz (*p_FDR_ <* 0.05) at T1, and only at 1 Hz (*p_FDR_ <* 0.01) at T2 (Figure 4D). A paired sample t-test (two tailed) showed no significant difference in RC1 *RSS* amplitude between T1 and T2 (*t*(1, 30) = 1.88*, p* = 0.07). In addition, a paired-sample t-test of projected amplitude at the first harmonic (significant at both testing time points) showed no significant difference between T1 and T2 (*t*(1, 30) = 0.33*, p* = 0.74). Phase comparisons between testing time points at 1 Hz (Figure 4E) were also not significant (circular t-test; *p* = 0.17).

**Figure 4:**
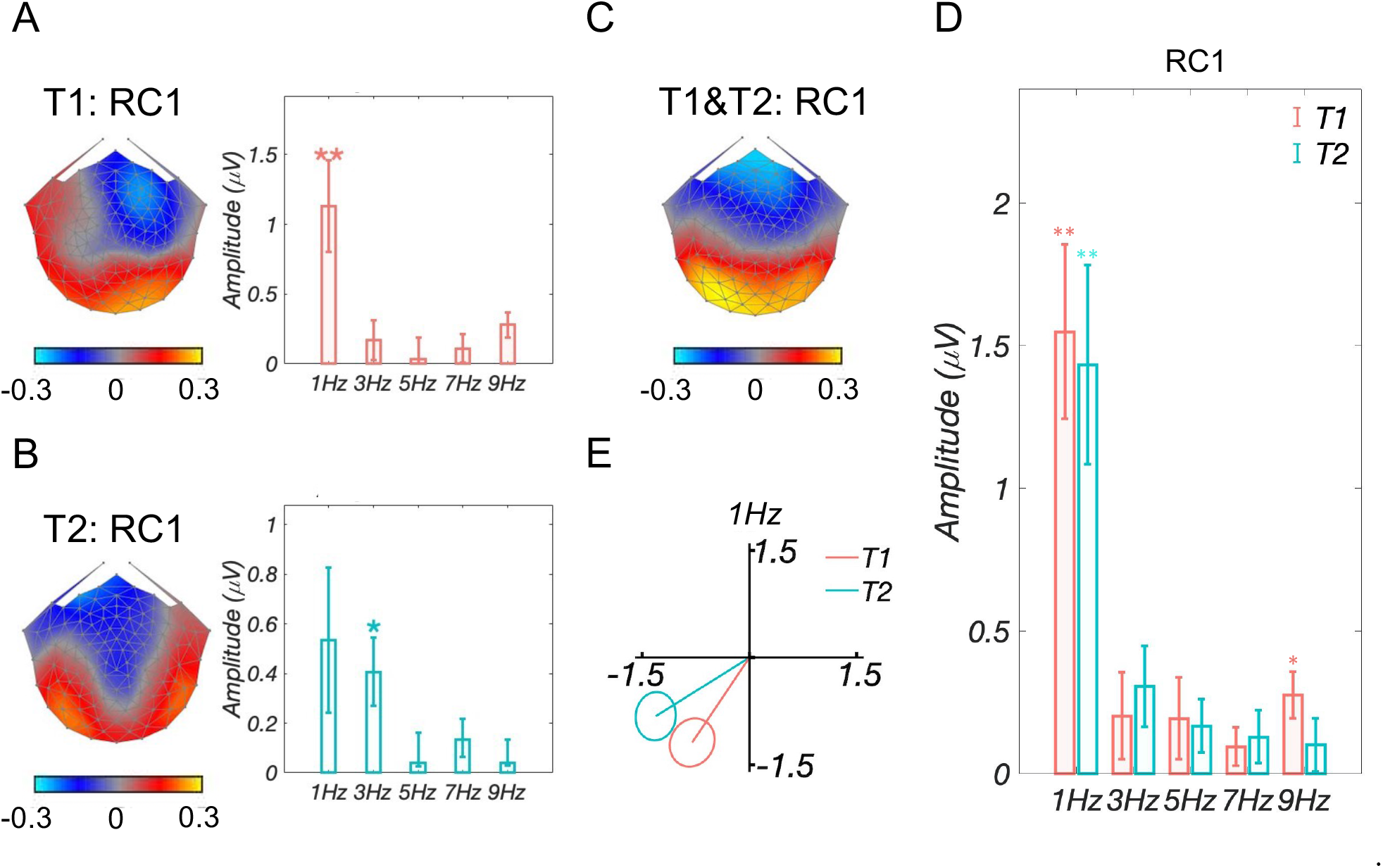
Alternate Analysis: Sensitivity to whole word representation (words–pseudowords) in early readers. A&B: Topographic visualizations of spatial filters and responses amplitudes for the maximally reliable component (RC1) trained on T1 and T2 separately; C: Topographic visualization of the spatial filter (more anterior left vOT) for RC1 trained on pooled T1 and T2 samples; D: Projected amplitude of RC1 (trained on T1 and T2 together) for each harmonic at T1 (orange) and T2 (blue), respectively. At T1, 1-Hz and 9-Hz harmonics were significant, while only the 1-Hz harmonic was significant at T2. There was no significant *RSS* amplitude difference (*p* = 0.07) between T1 and T2. Projected amplitude at the first harmonic showed no significant difference (*p* = 0.74) either between T1 and T2; E: Response data at the first harmonic (1 Hz, the only significant harmonic at both testing time points) presented in the complex plane, where amplitude information is represented as the length of the vector, and phase information in the angle of the vector relative to 0 degree (counterclockwise from 3 o’clock direction), ellipse indicates standard error of the mean (*SEM*). No significant phase difference (*p* = 0.17) was found between T1 and T2; *: *p_F DR_ <* 0.05, **: *p_F DR_ <* 0.01.

#### 3.1.3 Development of coarse print tuning (words–pseudofonts)

For responses to the words–pseudofonts contrast, aimed at coarse print tuning, the topography of RC1 at T1 includes bilateral peaks at posterior vOT electrodes (Figure 5A). At T2, the topography is more lateralized and maximal at left vOT electrodes (Figure 5B & supplement Figure S1B). The projected data were statistically significant in the first three harmonics (1 Hz, 3 Hz, and 5 Hz, *p_FDR_ <* 0.001; corrected for 10 comparisons) at T1, and all five harmonics at T2 (1 Hz, 3 Hz, and 5 Hz, *p_FDR_ <* 0.001, 7 Hz and 9 Hz, *p_FDR_ <* 0.05; corrected for 10 comparisons, Figure 5C). A paired sample t-test (two-tailed) showed no significant difference in RC1 amplitude between T1 and T2 (*t*(1, 30) = 1.10*, p* = 0.28). Phase comparisons (Figure 5D) between testing time points at 1 Hz, 3 Hz and 5 Hz showed no significant difference (circular t-test; *p_FDR_ >* 0.37, corrected for 3 comparisons); phases were not compared for 7 Hz and 9 Hz given that signals were significant at only one testing time point.

**Figure 5:**
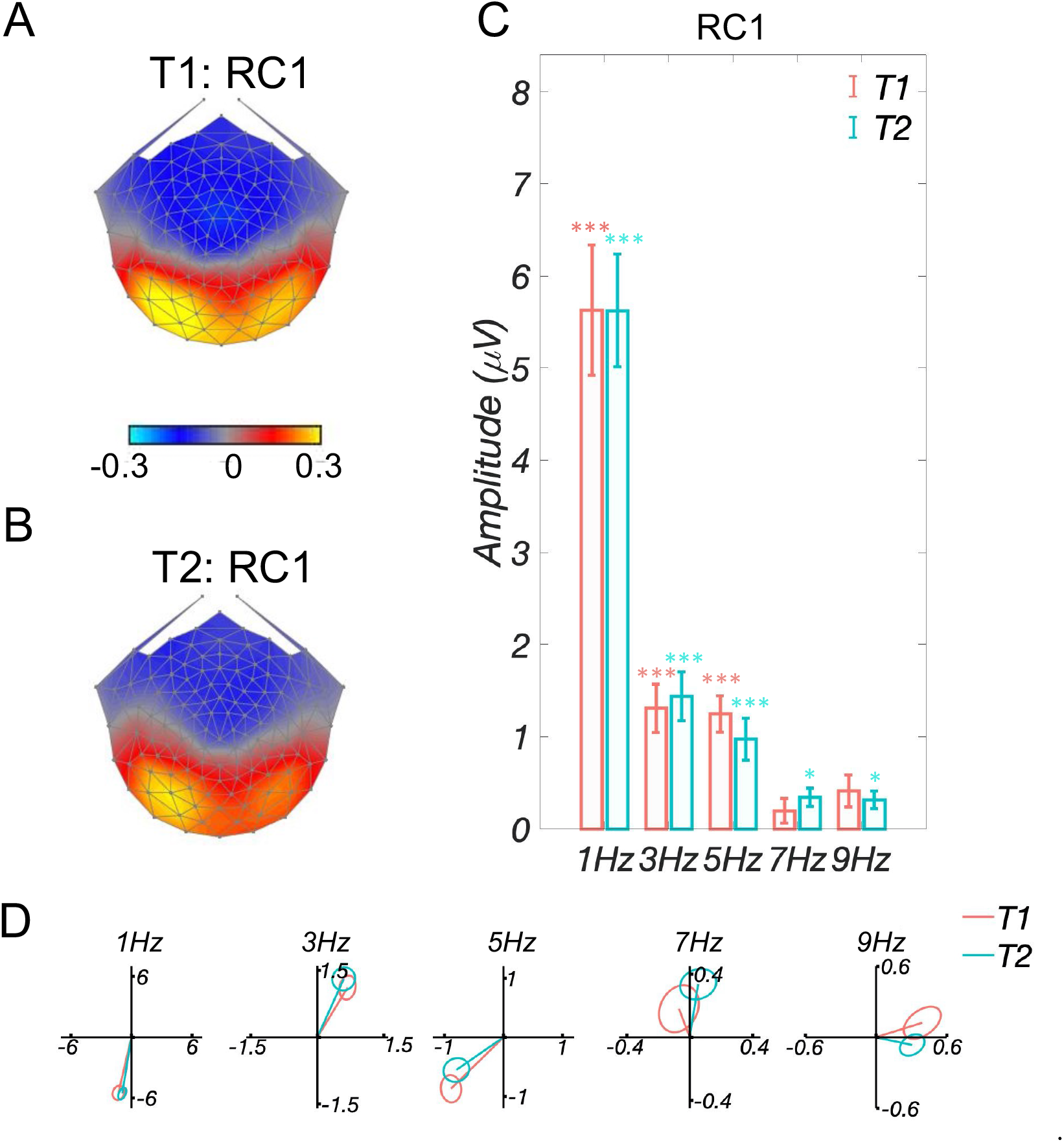
Alternate Analysis: development of coarse print tuning (words–pseudofonts) in early readers. A: Topographic visualization of the spatial filter (less left lateralized) for the maximally reliable component (RC1) at T1; B: Topographic visualization of the spatial filter (more left lateralized) for RC1 at T2; C: Projected amplitude of RC1 for each harmonic at T1 (orange) and T2 (blue), respectively. The first three harmonics were significant at T1, and all five harmonics were significant at T2. *RSS* amplitude did not differ significantly (*p* = 0.28) between T1 and T2. D: Response data in the complex plane, with T1 in orange and T2 in blue. Amplitude (vector length) and phase (vector angle, counterclockwise from 0 degrees at 3 o’clock direction) overlap between two testing time points especially for significant harmonics (first three harmonics). Ellipses indicate standard error of the mean. No significant phase differences (1 Hz: *p_F DR_* = 0.61, 3 Hz: *p_F DR_* = 0.61, 5 Hz: *p_F DR_* = 0.61) were found at first three significant harmonics between T1 and T2. *: *p_F DR_ <* 0.05, ***: *p_F DR_ <* 0.001.

### 3.2 Brain-behavior relationships

Brain-behavior analyses were performed to assess the relationship between component-space EEG amplitudes and behavioral reading abilities: Rapid naming abilities (RANcolor, RANletter), word reading efficiency (TOWRE), and word decoding ability (WJ). See *Behavioral Assessments* in *Materials and Methods* section and Table 1 for detailed information.

At T1, no clear relationships were found between response amplitudes of RC1 and reading scores for all three contrasts either before or after outlier removal (all *p_FDR_ >* 0.17, corrected for 4 comparisons).

However, brain-behavior relationships changed with 2 years of reading instruction in school (T2), as illustrated by significant linear relationships between response amplitudes to words–pseudofonts with WJ (word decoding, *r* = 0.5*, p_FDR_ <* 0.05, corrected for 4 comparisons, Figure 6A), as well as pseudowords–nonwords with RANletter (*r* = 0.4*, p_FDR_ <* 0.05, corrected for 4 comparisons) and WJ (word decoding, *r* = 0.7*, p_FDR_ <* 0.001, corrected for 4 comparisons, Figure 6B), after removal of outliers. No significant relationships were found between response amplitudes to the words–pseudowords contrast and reading scores (all *p_FDR_ >* 0.28).

**Figure 6:**
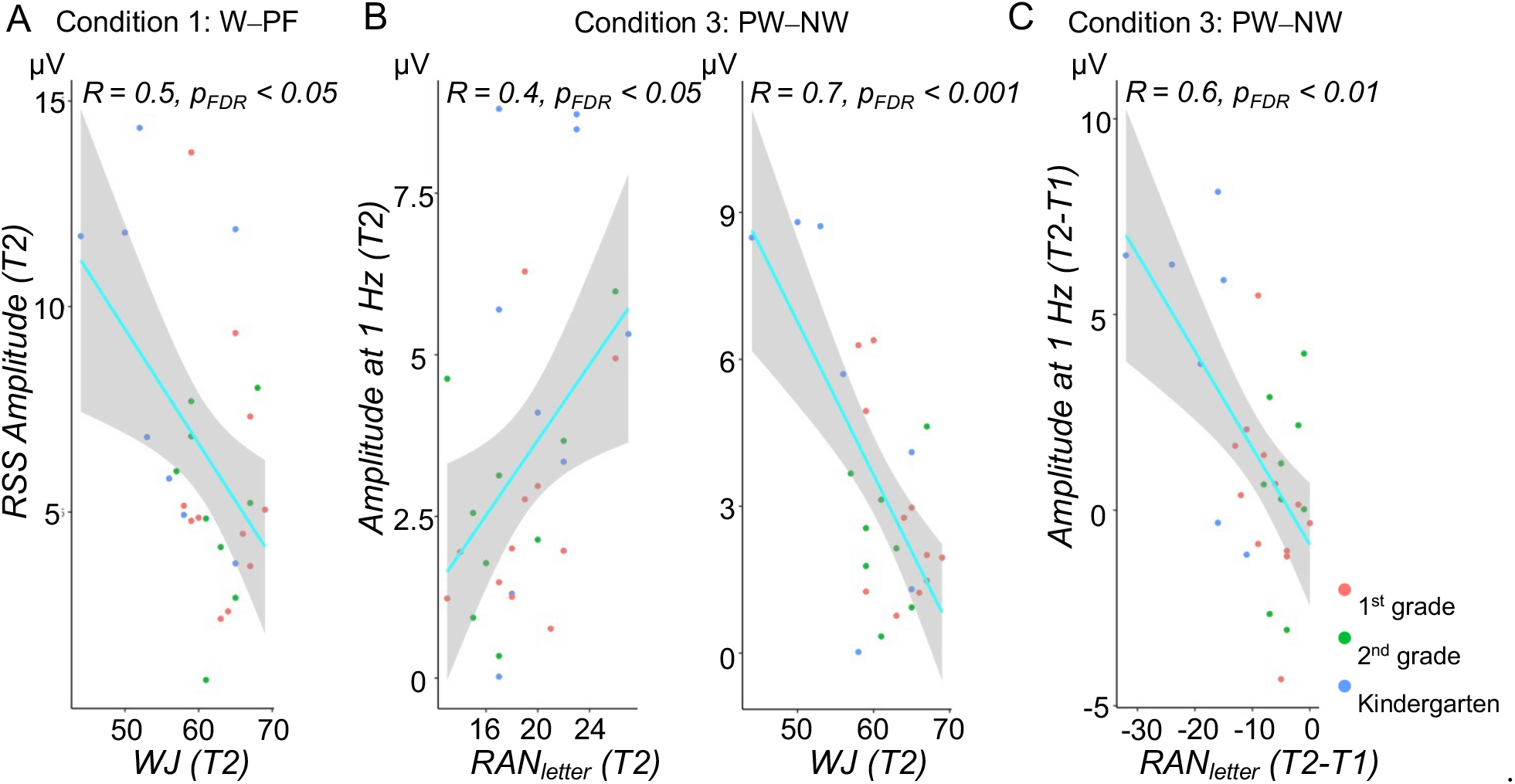
Statistically significant correlations between reading and brain response amplitudes for words–pseudofonts and pseudowords–nonwords, after outlier removal. A: Regression between *RSS* amplitude for words–pseudofonts and word decoding abiliy (Woodcock-Johnson, WJ) at T2; B: Regressions between response amplitude at 1 Hz for pseudowords–nonwords and rapid automatized naming of letters (RANletter) and WJ at T2; C: Regression between response amplitude improvement (Amplitude at 1 Hz*_T_* _2_ Amplitude at 1 Hz*_T_* _1_) at 1 Hz of pseudowords-nonwords and RANletter improvement (RANletter*_T_*_2_ RANletter*_T_* _1_). No significant relations were found for words–pseudowords. Note: The larger value of WJ and the smaller value of RANletter represent better reading skill.

Even more interestingly, we found a strong correlation between brain signal *improvement* (Amplitude at 1 Hz*_T_*_2_ *−* Amplitude at 1 Hz*_T_*_1_) for visual word form structure (pseudowords-nonwords) and letter reading speed *improvement* (RANletter*_T_*_2_*−*RANletter*_T_*_1_, *r* = 0.6*, p_FDR_ <* 0.01, after outlier removal and correction for five comparisons), see Figure 6C ^2^.

#### 3.2.1 Base RCA Results

We performed RCA on EEG responses at the base frequency and its harmonics (2 Hz, 4 Hz, 6 Hz, 8 Hz, and 10 Hz) in order to investigate neural activity underlying low-level visual processing. Similar with alternate results, we focused the base results on the maximally correlated component RC1. Figure 7A displays the topographies of RC1 separately for T1 (top) and T2 (bottom). RC1s are similar across testing time points, which are distributed over bilateral occipito-temporal region. Amplitudes (bar plots) are presented in Figure 7B. Figure 7C presents projected data (i.e., projecting data through the spatial filter) in the complex plane and shows overlapping amplitudes (vector lengths) and phases (vector angles) across three conditions and time points. Two-way repeated-measures ANOVA with within-subjects factors condition and testing time point revealed that neither the main effects nor the interaction were significant (all *F <* 2.61, all *p >* 0.1) on the projected amplitudes at each harmonic for RC1. Comparisons of phase across three conditions within a testing time point showed no significant difference (circular ANOVA, T1: all *F <* 2.61, all *p >* 0.1, T2: all *F <* 2.61, all *p >* 0.1). Thus, we consider the responses at the base frequency to be comparable across conditions and testing time points.

**Figure 7:**
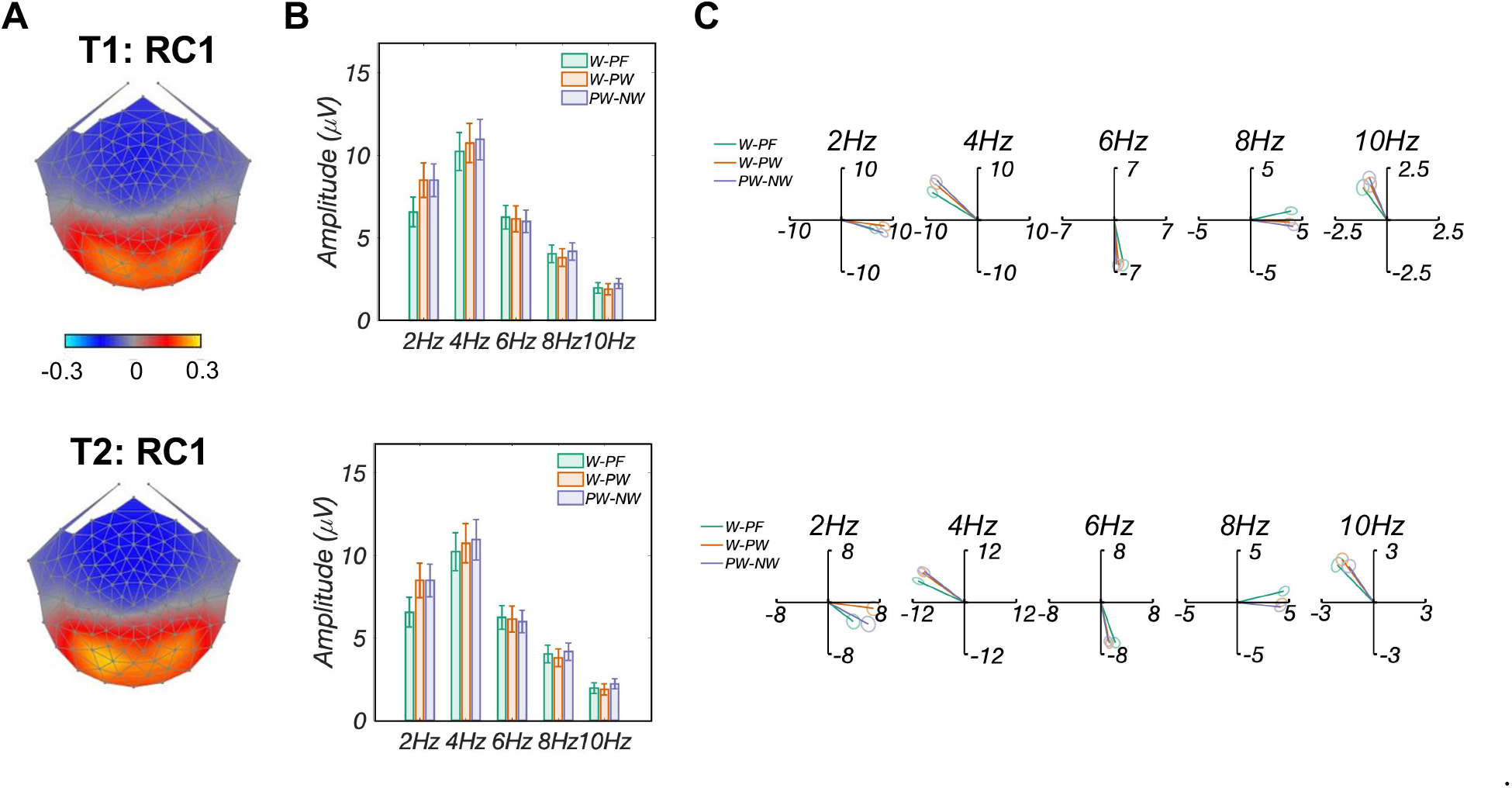
Base analysis comparisons across three conditions and two testing time points. A: Topographic visualizations of the spatial filters (T1: top; T2: bottom) for the first component (RC1); B: Comparison of projected amplitude across three conditions (T1: top; T2: bottom). Response amplitudes across five harmonics did not differ significantly across the three conditions and two testing time points (*p >* 0.1); C: Response data of three conditions presented in the 2D complex plane (T1: top; T2: bottom), where amplitude information is represented as the length of the vectors, and phase information in the angle of the vector relative to 0 degrees (counterclockwise from 3 o’clock direction), ellipse indicates standard error of the mean (*SEM*). Results showed overlapping amplitudes (vector lengths) and phases (vector angles) across three conditions and time points

### 3.3 Behavioral Results

We computed *d’* based on the z-transformed probabilities (the mean and standard deviation (SD) across three conditions are summarized in Table 2.) of hits and false alarms (Macmillan & Creelman, 2004). A two-way repeated-measures ANOVA of *d’* revealed a main effect of testing time point, with a higher hit rate at T2 compared to T1 (*F* (1, 185) = 17.04*, p <* 0.001). No significant difference was found across the three conditions (*F* (2, 185) = 1.95*, p* = 0.15), and no interaction between condition and testing time point was found (*F* (2, 185) = 1.80*, p* = 0.17).

**Table 2:** *d’* of the repetition detection task performance during EEG sessions for each of three conditions at T1 and T2. *d’* was computed based on the z-transformed probabilities of hits and false alarms. Values are mean(±SD).

|  | words–pseudofonts | words–pseudowords | pseudowords–nonwords |
| --- | --- | --- | --- |
| **T1** | 1.00($\pm 0.20$) | 1.04($\pm 0.30$) | 1.10($\pm 0.27$) |
| **T2** | 1.22($\pm 0.18$) | 1.53($\pm 0.24$) | 1.28($\pm 0.23$) |

## 4 Discussion

Over two years of school reading instruction, visual word form structure processing demonstrated striking patterns of growth: EEG responses nearly doubled in amplitude with increased left lateralization (Figure 3). More importantly, cortical entrainment of visual word form structure was significantly correlated with the growth in reading skill, especially in automatic rapid naming of letters. No such changes were observed for whole word representation processing, indicating that sensitivity to visual word form structure is uniquely linked to growth in reading. Theoretically, this finding sheds new light on current theories of word reading development and provides implications for models of visual word recognition. Practically, it yields insights for educational interventions and activities to improve reading fluency.

### 4.1 Development of brain sensitivity to visual word form structure (pseudowords–nonwords) in early readers

Behavioral studies have shown that sensitivity to visual word form structure develops rapidly through exposure to print (Deacon et al., 2013), even in fully unfamiliar and complex scripts (Chetail, 2017). Here, we measured changes in brain sensitivity (in terms of response amplitude and topography) underpinning word form structure processing over two years of reading instruction. The doubled response amplitude likely reflects increased sensitivity to orthographically reasonable word form structures, perhaps as they map to phonological patterns, as a result of reading expertise due to repeated exposure to print (Compton, 2000).

The maximally reliable spatial component of word form structure processing was distributed over the occipito-temporal (OT) regions and became increasingly left lateralized over two years (Figure 3A&B and supplement Figure S1). Presumptive activation over the left OT area here is consistent with previous studies showing its involvement in prelexical orthographic processing (Binder et al., 2006; Cohen et al., 2002; McCandliss & Noble, 2003). For example, an fMRI study with adult participants found sensitivity of the left lateral fusiform (occipitotemporal) gyrus to familiar over unfamiliar letter sequences (as measured by mean positional bigram frequency), even when these sequences did not resemble words (Binder et al., 2006). With the slower frequency rates (compared with 2 Hz oddball embedded in 10 Hz base in Lochy et al. (2016)) and a relatively explicit repetition detection task used in the present study, it is also likely that the brain responses to stimulus contrasts reflect orthographic and/or phonological decoding. Debska and colleagues (Dębska et al., 2019) suggested the involvement of automatic orthographic decoding in pseudoword recognition especially under orthographically demanding tasks (e.g., phoneme matching task in their study, repetition detection task in our study). Future studies with different tasks and/or more precise manipulations to control potential decoding are needed to improve our knowledge on this issue.

Previous studies have not examined the lateralization changes of word form structure processing in early readers. Instead, most previous findings are centered on the lateralization patterns of coarse neural tuning effect and support the assumption that the degree of left lateralization for print is dependent on the level of reading expertise with a script (Maurer et al., 2008; Spironelli & Angrilli, 2009). Indeed, we also found a left lateralization shift for coarse print tuning as in previous literature (Araújo, Faísca, et al., 2015; Brem et al., 2005; Hauk et al., 2006; Zhao et al., 2014). Moreover, the current study extends previous findings and additionally revealed a left lateralization shift for visual word form structure tuning. Right hemisphere activation is usually detectable in the early stages of learning to read, and tends to reduce or disappear as development proceeds (Turkeltaub et al., 2003). This reduced involvement of right hemisphere areas over the course of reading development reflects the experience-driven functional refinement of the skill zone for reading, which manifests as greater recruitment of left hemispheric OT regions (Turkeltaub et al., 2003). Of note, no lateralization changes were found for whole-word lexical processing (pseudowords–nonwords). The current findings on different patterns of lateralization changes for different levels of word information processing further reveal that such changes are not a genetic lateralization shift of left vOT over time. Rather, it shows specificity of letter and word form structure, at least during the early reading stage.

### 4.2 Correlations between brain sensitivity to visual word form structure and reading fluency

We also found that brain sensitivity to word form structure contributes to reading fluency. Children with larger response amplitudes to word form structure had better reading performance in rapid letter naming (RAN for letters) as well as word decoding (measured with letter-word identification using WJ) at T2 (Figure 6B). This finding aligns with the observation of brain sensitivity to visual word form structure in skilled adult readers but not in dyslexic individuals (Araújo, Faísca, et al., 2015). Such deviation between typical and dyslexic readers suggests a connection between sufficient tuning to visual word form structure and typical reading skills. In relation to this, a developmental ERP study demonstrated neural sensitivity to visual word form structure in 7-year-old children with high reading ability, but not in those with low reading skill (Zhao et al., 2014). Collectively, the current study and previous ERP studies provide evidence that the ability to detect and apply visual word form structure is a crucial factor promoting reading success (Conrad et al., 2013).

### 4.3 Correlations between brain sensitivity to coarse tuning and reading skill

Previous SSVEP studies (e.g., preschoolers in Lochy et al. (2016); first and second graders in van de Walle de Ghelcke et al. (2021)) revealed significant correlations between response amplitude for coarse print tuning and reading scores. In the current study, however, amplitude of the coarse tuning contrast was only related to reading skill (specifically word decoding ability) at T2 but not at T1. Such discrepancies may be due to the relatively explicit task (repetition detection task, rather than monitoring fixation color change in previous studies) and slower presentation rates in an alternation paradigm used in our design.

First, explicit tasks might focus purely on proficiency ability differences (Zobl, 1992), rather than implicit tasks that might ambiguously combine proficiency ability differences with spontaneous tendencies and strategies (Nosek et al., 2011). Children in a sample might have equal ability to carry out a computation when explicitly directed to do so (and thus no correlation with skill), yet still differ in their spontaneous tendency (Bennett-Branson & Craig, 1993) to carry out the task given an implicit task situation, and that implicit tendency may well correlate with skill level (Liang et al., 2021). In addition, we found the relationships between brain responses and reading skills grew stronger over 2 years via the more explicit task. This finding further supports the assumption that this is a longitudinal change in computational ability (evoked by an explicit task) rather than longitudinal changes in spontaneous tendencies (evoked by an implicit task). Hence, explicit tasks might hold advantages for tracking development in studies that focus more on ability differences when other influences (like spontaneous goal or strategy differences) are minimized.

Second, in contrast to previous fast frequency rates in an oddball paradigm (e.g., 2 Hz oddball and 10 Hz base in Lochy et al. (2016)), the slower rates (i.e., 1 Hz alternate and 2 Hz base) in a two-stimulus evenball design (e.g., alternating pseudowords and nonwords once per second) may also explain the discrepancy. Slower presentation rates may drive more attention to the visual word form encoding/decoding processes, which may enable the capture of processing stability of coarse tuning and result in diminished correlations between amplitude and reading skill, especially at the very beginning of reading acquisition (herein at T1).

Future studies outside of the scope of the current investigation will be explicitly designed to better capture the influences of different frequency rates and different task modulations on neural responses and brain-behavior correlations.

### 4.4 Correlation between development of brain signal and growth in reading fluency

Remarkably, our results showed that the development of brain sensitivity to word form structure is significantly correlated with growth in early reading skill. We found that early readers with larger changes in amplitudes to word form structure had higher improvement in RAN (letters not colors, Figure 6C).

A potential interpretation for this relationship is that beginning readers who are slow to identify individual letters may not activate the letters in memory close enough in time to encode the letter combinations that occur most frequently in print (Bowers et al., 1994). This is consistent with the assumption that the inability to sufficiently automatize letter recognition interferes with letter string processing and growth of orthographic knowledge (Manis et al., 2000). For instance, Bowers and colleagues reported a deficit in word form structure learning in their RAN deficit group (Bowers et al., 1999). Numerous studies have suggested that RAN is a strong predictor of reading fluency (for a meta-analysis, see Araújo, Reis, et al. (2015); for a review, see Georgiou et al. (2013)). Given that no significant correlations were found with RAN for colors, the current study provides support for the idea that the growth of brain sensitivity to visual word form structure specifically relates to reading fluency and reading development, instead of general articulation speed.

In addition, it might be also possible that the development of brain sensitivity to word form structure reflects improvement in grapheme-to-phoneme conversion and/or decoding. During early reading acquisition, children first need to learn to decode letters into words by identifying letters and mapping letters to corresponding sounds. Years later, a form of perceptual expertise emerges in which groups of letters are rapidly and effortlessly conjoined into integrated visual percepts (McCandliss et al., 2003). Supporting this, researchers have also found automaticity of word decoding to be a critical component of fluent reading ability (usually measured with RAN as in the present study) and is essential for high levels of reading achievement (Pikulski & Chard, 2005; Roembke et al., 2019). Moreover, slower presentation rates and the 3-in-a-row repetition detection task in the current study might draw more cognitive resources and attention to the decoding process.

To conclude, the development of visual word form structure encoding and decoding mechanisms is an important aspect of word recognition skill that allows readers to process letters and letter combinations rapidly and fluently. It will be interesting to explore the similarities/differences between effects of encoding and decoding on visual word form structure learning in future studies.

### 4.5 Evidence for sensitivity to whole word representation in early readers

Brain responses to word-level representations have also been studied previously; inconsistent results are presumably due to this effect being less pronounced and more task- and developmental stage-dependent (Maurer et al., 2006; Xue et al., 2008). For example, past ERP studies (Zhao et al., 2014; Maurer et al., 2006) reported larger amplitudes for words than for pseudowords in children (7-8 y old 2nd graders), which was also demonstrated in adult ERP (Hauk et al., 2012) and SSVEP (Lochy et al., 2015) studies. In contrast, Brem et al. (an ERP-fMRI study with 10 y old children and adolescents, Brem et al. (2009)) and Hauk et al. (an ERP study with adults, Hauk et al. (2006)) reported stronger neural responses for pseudowords than for words. Yet other studies, including ERP (9-13 y old pre-adolescents in Araújo et al. (2012); 7.6 y old second graders in Eberhard-Moscicka et al. (2015); 6.5 y old kindergarteners in Maurer et al. (2006)) and SSVEP (5 y old preschoolers in Lochy et al. (2016); 6-7 y old first and second graders in van de Walle de Ghelcke et al. (2021)) studies found null effects.

The current study was able to capture brain signals related to representation of familiar words over occipital and temporal regions (Figure 4), potentially due to slower presentation rates in an SSVEP alternation paradigm with a repetition detection task. However, no correlations between response amplitudes and reading skills were found for this contrast in either of the two testing time points. The ambiguity of brain-behavioral relationships here might be due to the small size of this word representation effect (Eberhard-Moscicka et al., 2015). Future studies can examine this issue further by recruiting participants spanning a larger range of word representation processing and/or by employing more explicit tasks (e.g., lexical decision task), which might better engage this type of processing. An alternative explanation is that the words we used were all very high frequency, simple, and short (3 letters). Such stimuli do not require much attention on word decoding, resulting in less robust correlations with reading skills related to decoding (WJ) and automatized letter naming (RAN for letters). This speculation could be resolved through a further test of word form structure learning and reading development with more complex, longer stimuli with variable frequencies.

### 4.6 Implications for reading models and educational neuroscience research

Response amplitudes to coarse neural tuning (words–pseudofonts) were significantly correlated with amplitudes to word form structure (pseudowords–nonwords), instead of whole word representation (words–pseudowords), at both testing time points (see supplement, Figure S3). Moreover, similar correlations between brain signal and reading skill (mainly word decoding ability) were found for contrasts of coarse neural tuning and word form structure. These findings may indicate that, at least during early reading acquisition, the words-pseudofonts contrast often used in previous studies might primarily pick up information about sublexical letter form and/or word form structure rather than whole-word lexical processing.

Despite high correlation and information overlap between coarse tuning (words–pseudofonts) and word form structure tuning (pseudowords–nonwords), developmental changes—in terms of response amplitude—occurred only for word form tuning and not coarse tuning. Several factors may play a role in the disparity. First, cognitive and attentional processes of visual word form structures in very high-frequency familiar words and unfamiliar pseudowords might be different. Compared with very familiar 3-letter words which most likely can be automatically encoded (Spironelli & Angrilli, 2009), unfamiliar pseudowords possibly require additional attention and processing energy (e.g., decoding) due to lack of visual familiarity (Maurer et al., 2005). This increase in attention and potential effortful decoding might explain the amplitude increase mostly on these unfamiliar pseudowords forms. Second, it is likely that developmental changes of coarse tuning happen earlier than sublexical visual word form structure tuning. A previous SSVEP study found automatic encoding of familiar words (vs. pseudofonts) in preschoolers (Lochy et al., 2016). In an ERP study, Maurer and colleagues found increased coarse tuning in second graders compared with non-reading kindergarteners (Maurer et al., 2006). Our samples, however, included children spanning kindergarten to second grade with a two-year follow up, which might have missed the key increasing stage (i.e., between kindergarten to second grade). Thus, longitudinal studies with only beginning readers (i.e., kindergarteners) might better capture the developmental changes of coarse tuning. Finally, distinguishing words from pseudofonts involves different demands from distinguishing pseudowords from nonwords. Future studies comparing words and pseudowords processing in the same base context (e.g., pseudofonts) would speak directly to this question.

Nevertheless, our findings of developmental changes of visual word form structure tuning over two years and the relationship with reading skill (mainly decoding) improvement suggest that word form structure processing plays a crucial role in reading fluency and reading growth in early readers.

Sensitivity to word form structure has been rarely or not even taken into account in current models of visual word recognition (Chetail, 2015, 2017; Harm & Seidenberg, 2004). Most of these models, including Dual Route Cascaded (DRC) model (Coltheart et al., 2001) and triangle model (Harm & Seidenberg, 1999), have focused mainly on properties of lexical-semantic and phonological representations and processes. It will be important for future research and reading models to implement visual word form structure in a manner analogous to what has been done with semantics and phonology. This extra word form structure component of the model would provide insights for bridging educational practice and neuroscience to improve early reading fluency.

### 4.7 Limitations

Several limitations of the present study should be addressed. First, we examined the longitudinal changes of the hierarchical processing of word information in an unbalanced condition order. We started with the intention to replicate and extend the coarse print tuning effect (words–pseudofonts) that has been investigated in numerous studies. Then, we aimed to functionally dissociate two related functions of the visual word form system linked to two orthographic grain sizes (lexical and sublexical) by contrasts of words–pseudowords and pseudowords–nonwords, respectively. As a consequence, we cannot rule out the possibility of serial order carryover effects from one condition to the next. Future studies may use counterbalanced condition orders to more precisely capture the condition effects, although this would come at the cost of a much larger sample size. Second, the small sample sizes of the grade groups limit us from making conclusions about different brain-behavior relations based on different grades. Future studies with larger sample sizes in each age and/or grade group will enhance our ability to trace the developmental trajectories of brain-behavior relationships in children at different stages of learning to read. Finally, an explicit task was used in the current study, which was revealed might hold advantages for tracking development in studies that focus more on reading ability differences (see Discussion). However, the explicit task increased the difficulty of disentangling encoding and decoding processing of words. Future studies including both implicit and explicit tasks may help to clarify whether the (development) changes of visual word form structure processing are specific to encoding or decoding, or even both. Nevertheless, we believe our results still provide interesting insights into the developmental profile of different levels of word information and their effects on reading skill.

## 5 Conclusion

This longitudinal study demonstrates developmental changes—in terms of response amplitudes and left lateralization—of visual word form structure processing in early readers. Moreover, the word form structure effect became stronger in faster readers, supporting its functional role in early reading ability. No such changes were observed for responses to whole word representation. Taken together, our results suggest that word form structure processing, which is indispensable for further accessing the phonology and semantics of written words, may be an important factor influencing reading ability growth. Such knowledge is crucial to better understand how children develop sensitivity to visual words and is insightful for models of visual word recognition.

## Data Availability Statement

The data sets generated for this study are available on request to the corresponding author.

## Author Contributions

F.W. and B.D.M. conceived the study. F.W., S.M., R. S. G., E.Y.T, L.R.H., and Q.T.H.N. conducted the experiment(s). F.W. and B.K. analyzed the data. F.W. wrote the original draft, B.K., L.R.H., E.Y.T., R.S.G., A.M.N., and B.D.M. edited the draft. All authors reviewed the manuscript.

## Ethics Approval Statement

The study was approved by the Institutional Review Board of Stanford University. A parent or legal guardian of each participant received a written description of the study and gave written informed consent before the session; each participant also assented to participating.

## Permission To Reproduce Material From Other Sources

Figures were created in MATLAB (2020a) using publicly available code (https://github.com/svndl/rcaExtra)

## Acknowledgments

We thank the students, their families, and teachers for participating. We also thank Rachana Pillai for her help with data collection.

## Abbreviations

RCA: Reliable Components Analysis
SSVEP: steady-state visual evoked potentials

## Footnotes

1 At T1, a total of 57 healthy, English-speaking children participated. Out of these initial 57 subjects, nine were excluded from further analyses (e.g.,EEG data quality issues). The remaining sample (N=48) included 15 kindergarteners, 16 first graders and 17 second graders (see Wang et al. (2022)).

2 Regression models in Figure 6 still hold after running models with age as a covariate.

