## Supplement Text for "Progress in elementary school reading linked to growth of cortical responses to familiar letter combinations within visual word forms"

### 1 Supplementary Material

#### 1.1 SI Method

##### 1.1.1 Analysis of the correlations of alternate amplitudes across three conditions

We examined amplitude relationships across three conditions using $RSS$ projected amplitude from alternate RCA output for words–pseudofons and 1 Hz (the only significant harmonic) amplitude for words–pseudowords and pseudowords–nonwords. To test the association between amplitudes for different conditions at different developmental stages, we performed linear regressions of response amplitudes, separately for T1 and T2. Outliers determined by Cook's Distance Cook (1977) based on the regression model were removed if they exceeded the $4/n$ threshold ($n$ total data points).

##### 1.1.2 Analysis of lateralization of brain responses to three conditions

To investigate lateralization of brain responses to each stimulus contrast, first, we simulated regions of interest (ROIs) for the left and right visual word form area (VWFA), which are specialized for computation of abstract visual word forms Cohen et al. (2000) using EEG source simulation toolbox Barzegaran et al. (2019). Second, we projected electrode-by-frequency-by-trial sensor space data matrix at both testing time points onto left and right VWFA ROIs, respectively, resulting projected data matrix of component-by-frequency-by-trial. Finally, a three-way repeated-measures ANOVA with within-subjects factors of condition, testing time point, and hemisphere was computed on response amplitude (Figure S1A).

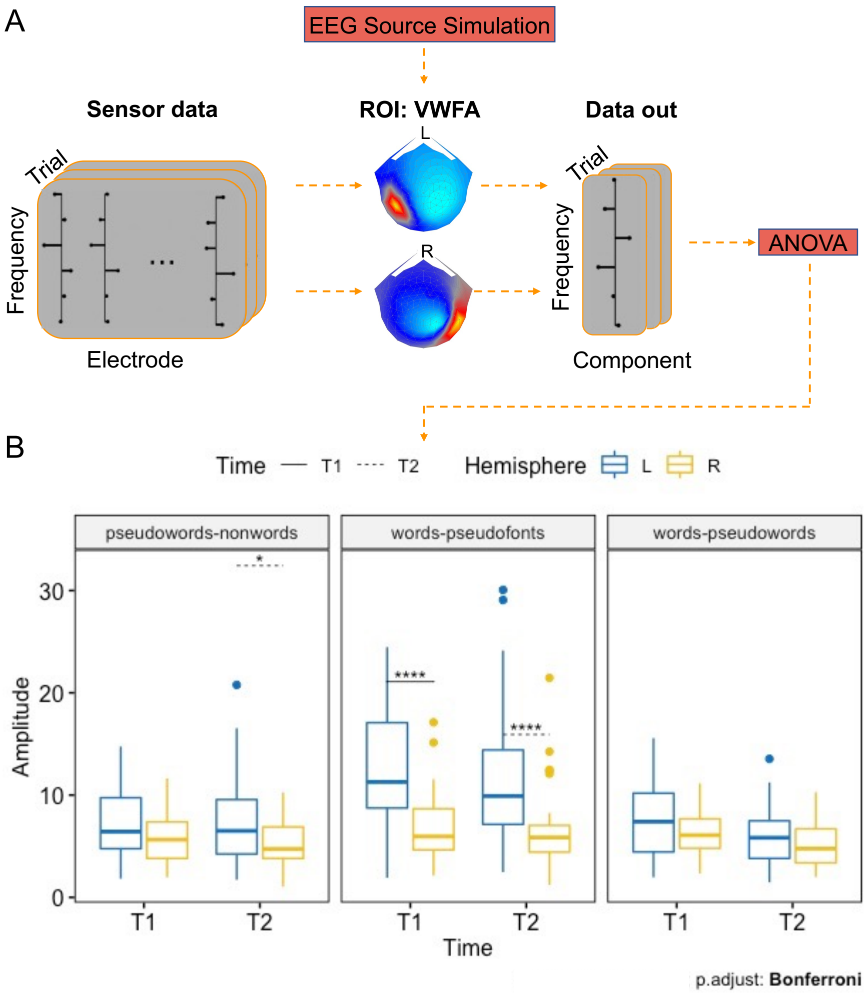


Figure S1: **Procedure and test results for lateralization of brain responses in three conditions.** A: Procedure of testing brain responses lateralization. First, regions of interest (ROIs) for left and right visual word form area (VWFA) were simulated using EEG source simulation toolbox (Barzegaran et al., 2019). Second, we projected electrode-by-frequency-by-trial sensor data matrix at both testing time points onto left and right VWFA ROIs, respectively, in which the resulting output data (i.e., projected data) turned to a matrix of component-by-frequency-by-trial. Finally, projected data were analyzed with a three-way repeated-measures ANOVA with within-subject factors condition, testing time point, and hemisphere; B: Results of lateralization test. Significant main effects of condition ($p < 0.001$) and hemisphere ($p < 0.001$), as well as interaction effect of condition by hemisphere ($p < 0.001$) were revealed. Post-hoc tests showed that brain responses were left lateralized for coarse tuning effect (words-pseudofonts) at both T1 and T2; for visual word form structure processing (pseudowords-nonwords), brain responses were more left lateralized at T2 compared with T1. Trend but not statistically significant left lateralization effects were found for whole word presentation processing. *: $p_{adj} < 0.05$; ***:$p_{adj} < 0.001$

#### 1.2 SI Results

##### 1.2.1 Permutation Test Results

Permutation testing was performed on alternate and base RCA coefficients. As shown in Figure S2, for alternate RCA calculations, permutation testing revealed statistically significant (permutation test $p_{FDR} < 0.001$, each corrected for 5 comparisons) RC1–5 coefficients for words–pseudofonts contrast at both T1 (Figure S2A) and T2 (Figure S2B). For words–pseudowords contrast, RC1–2 and RC1–5 were statistically significant (permutation test $p_{FDR} < 0.001$, each corrected for 5 comparisons) at T1 and T2, respectively. For pseudowords–nonwords, RC1 and RC1-3 were statistically significant (permutation test $p_{FDR} < 0.05$, each corrected for 5 comparisons) at T1 and T2, respectively. For base RCA calculations, RC1–5 coefficients were all statistically significant (permutation test $p_{FDR} < 0.001$, each corrected for 5 comparisons) at both T1 and T2.

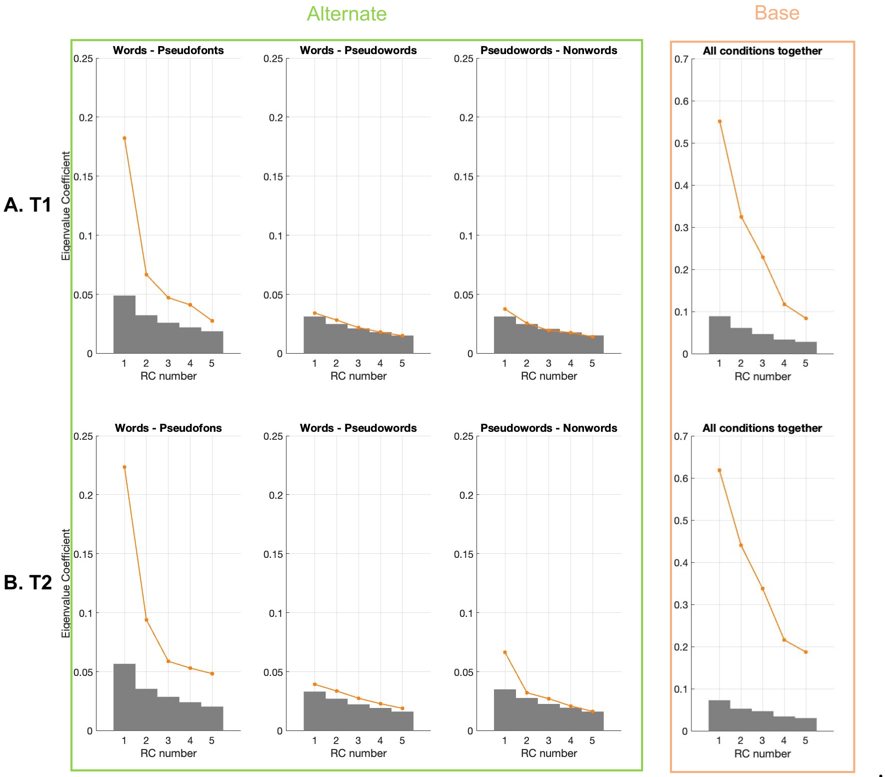


Figure S2: **Statistical analyses of eigenvalue coefficients of RC1–5 at (A) T1 and (B) T2.** The line plots denote the distribution of component coefficients for each RCA calculation, and the shaded gray area denotes the 95th percentile of each component's null distribution. Alternate: RC1–5 coefficients for words–pseudofonts were all statistically significant (permutation test $p_{FDR} < 0.001$, each corrected for 5 comparisons) at both T1 and T2; for words–pseudowords, RC1–2 were statistically significant (permutation test $p_{FDR} < 0.05$, corrected for 5 comparisons) at T1, while RC1–5 were all statistically significant (permutation test $p_{FDR} < 0.01$, corrected for 5 comparisons) at T2; for pseudowords–nonwords, RC1 and RC1-3 were statistically significant (permutation test $p_{FDR} < 0.05$, each corrected for 5 comparisons) at T1 and T2, respectively. Base: RC1–5 coefficients for base RCA performed on three conditions together were all statistically significant (permutation test $p_{FDR} < 0.001$, each corrected for 5 comparisons) at both T1 and T2.

##### 1.2.2 Correlations between amplitudes of three conditions

Linear regressions between alternate amplitudes of three conditions were performed to assess the relationship between different levels of word information processing. As shown in Figure S3, response amplitudes for words–pseudofonts contrast (i.e., coarse neural tuning) were significantly correlated with amplitudes for pseudowords–nonwords contrast (i.e., sensitivity to word form structure), instead of words-pseudowords contrast (i.e., sensitivity to whole word representation), at both T1 (Figure S3A, $p_{FDR} < 0.01$, corrected for 3 comparisons) and T2 (Figure S3B, $p_{FDR} < 0.001$, corrected for 3 comparisons). No significant relations were found between response amplitudes to words-pseudowords contrast and pseudowords–nonwords contrast.

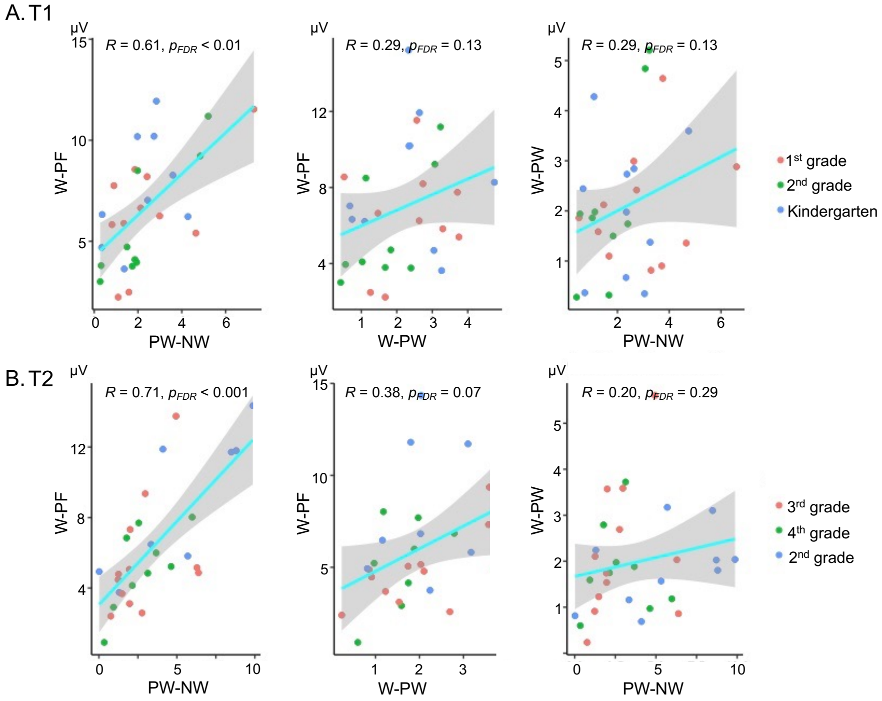


Figure S3: **Correlations between amplitudes of the three conditions at two testing time points.** Linear regressions between amplitudes for words–pseudofonts (W–PF) and pseudowords–nonwords (PW–NW), words–pseudofonts (W–PF) and words-pseudowords (W–PW), and words-pseudowords (W–PW) and pseudowords–nonwords (PW–NW) at T1 (A) and T2 (B). Left panel: Statistically significant correlations between response amplitudes for words–pseudofonts (W–PF) contrast (i.e., coarse neural tuning) and pseudowords–nonwords (PW–NW) contrast (i.e., sensitivity to word form structure) at both T1 ($p_{FDR} < 0.01$, corrected for 3 comparisons) and T2 ($p_{FDR} < 0.001$, corrected for 3 comparisons); middle panel: Response amplitudes for words–pseudofonts (W–PF) contrast (i.e., coarse neural tuning) were not significantly correlated with amplitudes for words-pseudowords (W–PW) contrast (i.e., sensitivity to whole word representation). right panel: No significant relations were found between response amplitudes to words-pseudowords (W–PW) contrast and pseudowords–nonwords (PW–NW) contrast.

##### 1.2.3 Lateralization of brain responses to three conditions

A three-way repeated-measures ANOVA with within-subjects factors condition, testing time point, and hemisphere on $RSS$ projected amplitudes of three stimulus contrasts was computed to test the lateralization changes of brain responses. As shown in Figure S1B, we found statistically significant main effects of condition ($F(2, 60) = 30.25, p < 0.001$) and hemisphere ($F(1, 30) = 36.28, p < 0.001$), as well as an interaction effect between condition and hemisphere ($F(2, 60) = 30.19, p < 0.001$), on the RSS projected amplitude. Neither significant main effect of testing time point ($F(1, 30) = 1.57, p = 0.22$) nor significant interaction effects condition × testing time point ($F(2, 60) = 0.95, p = 0.39$), testing time point × hemisphere ($F(2, 60) < 0.1, p = 0.99$), and condition × testing time point × hemisphere ($F(2, 60) = 0.38, p = 0.69$) were found. Post-hoc t tests showed that brain responses to coarse print tuning (words–pseudofonts) were left lateralized at both testing time point (both $p < 0.001$, Bonferoni corrected). Brain responses to visual word form structure processing (pseudowords–nonwords) were more left lateralized at T2 ($p < 0.001$, Bonferroni corrected) than T1 ($p = 0.168$). Trend but not statistically significant left lateralization effects were found for brain responses to whole word representation processing (words–pseudowords).
